# Multi-level repression ensures tight translation control of C9ORF72

**DOI:** 10.1101/2025.08.09.669471

**Authors:** Solomon A. Haizel, Alfredo Vidal Ceballos, Alice Nayoung Ko, Lucas Lyu, Lauren Manning, Xinyi Ge, Violet Park, Huijing Zou, Churong Cheng, Richmond Afful, Serena Kher, Anthony Widjaja, Joanetta Jean-Baptiste, Sasha Khazhinsky, Pieter Spealman, Alisha N. Jones, Leos Shivaya Valasek, Christine Vogel

## Abstract

C9ORF72 is involved in multiple neuronal functions and a major factor in Amyotrophic Lateral Sclerosis, a fatal neurodegenerative disease. Pathogenicity is implemented through an intronic repeat expansion in the gene’s mRNA leader that produces long, repetitive RNA and dipeptides when translated through repeat associated non-canonical translation. The expanded repeat also suppresses transcription of the major transcript variant (Variant 2) which comprises 90% of the C9ORF72 levels in healthy cells, leading to an additional loss-of-function effect. Examining the sequence, structure, and ribosome occupancy of Variant 2’s mRNA leader in detail, we found that translation of C9ORF72 is tightly regulated through a multi-layered system. To deconvolute this system, we conducted a mutant screen and developed a method to evaluate the results quantitatively. We found that near-cognate start codons in all three frames, a start-stop element in frame 2, a uTranslon in frame 1, and secondary structures cooperate to keep baseline C9ORF72 translation low. Further, initiation at upstream near-cognate start codons reduces the impact of both the start-stop and the uTranslon, effectively buffering their repressive function and creating a feedforward loop. A strong secondary structure in the Variant 2-specific sequence further modulates initiation at all the elements. We hypothesize that this buffered repressor system has evolved to ensure reliable, noise-insensitive expression of this critical regulator of neuronal function. Finally, we used ADAR-based RNA editing of specific sites in Variant 2’s mRNA leader, increasing translation of the gene and providing a novel avenue to rescue the loss-of-function phenotype in disease.

## Introduction

C9ORF72 is a guanine nucleotide exchange factor involved in many processes that underpin neuronal function and survival. It is highly expressed in the brain, primarily in the cerebellum ^1^. It plays an essential role in autophagy, facilitating the degradation of damaged organelles and protein aggregates through its interaction with autophagy-related proteins such as ULK1 and RAB proteins ^2–5^. Additionally, C9ORF72 is integral to endosome-lysosome function, contributing to membrane trafficking and the clearance of cellular waste by modulating the activity of RAB GTPases, which are vital for vesicle transport and fusion ^6^. Beyond these, emerging evidence suggests C9ORF72 participates in synaptic vesicle trafficking, supporting neurotransmitter release and neuronal communication ^2–4,7^.

The C9ORF72 gene produces three major transcript variants that differ in the inclusion of a non-coding exon present in the 5’ UTR ^1^. The variants have been named Variant 1, 2, and 3, referring to the Refseq sequences NM_145005, NM_018325, NM_001256054, respectively ^1^. Variants 1 and 3 feature a short and long version of exon 1a in their 5’untranslated region (UTR), respectively; while Variant 2 carries exon 1b (**Figure 1a**). All three variants share exon 2 which contains the AUG start codon of the coding sequence (main translon). Variant 2 is the dominant transcript isoform contributing ∼70-100% of the total C9ORF72 transcript levels ^8^ and it is mainly expressed in the brain and spinal cord ^9^.

**Figure 1.**
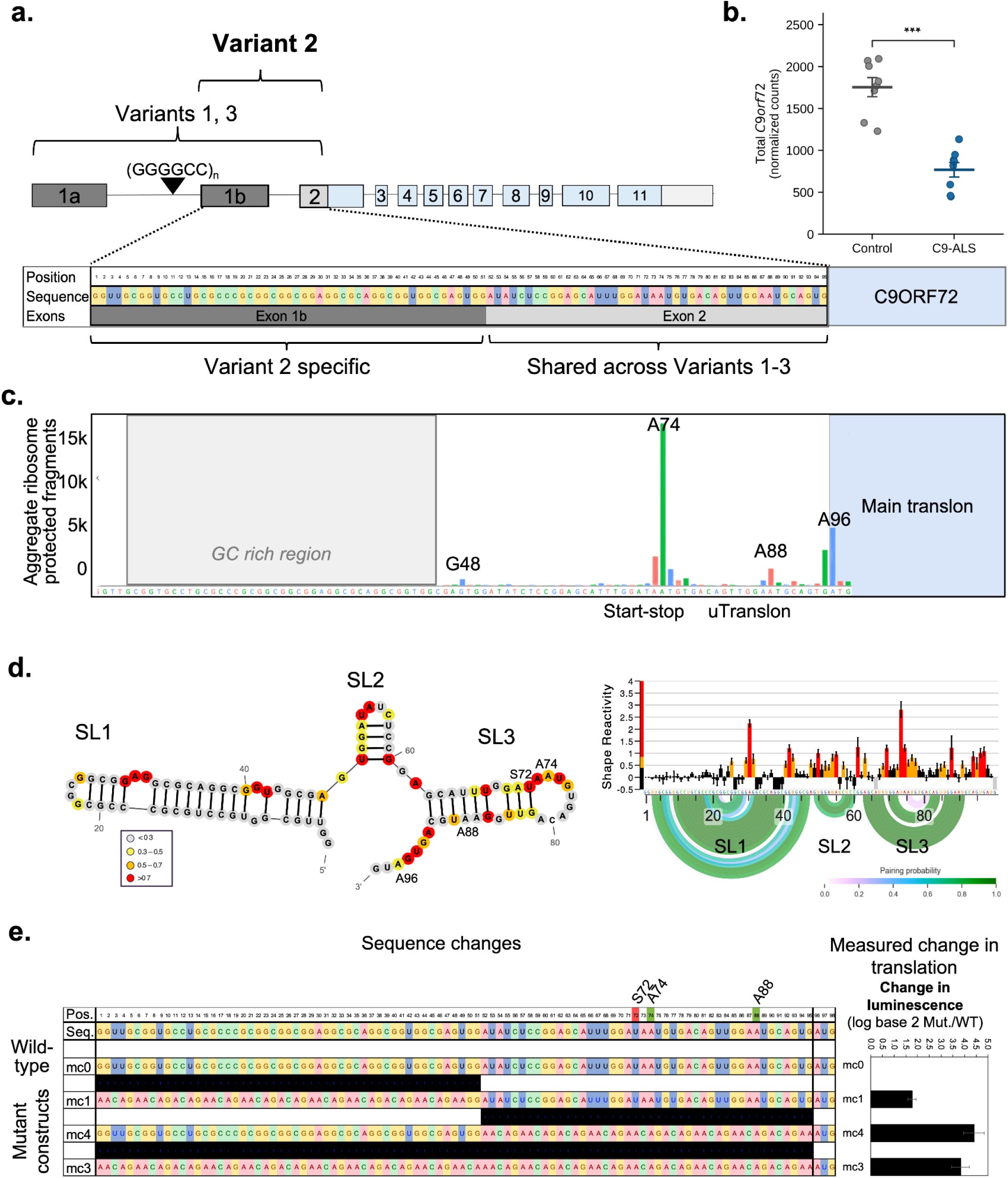
C9ORF72 Variant 2 is the dominant transcript isoform with multiple predicted translation regulatory elements. **a.** Schematic of the *C9ORF72* gene with the three known mature transcript variants. Exon 1a is shared by Variants 1 and 3, whereas exon 1b is specific to Variant 2 (V2). All three variants share exon 2, which contains the AUG start codon of the coding sequence. The black triangle indicates intronic (GGGGCC)*n* hexanucleotide repeat between exons 1a and 1b, pathogenically expanded in C9-ALS/FTD. The 95-nucleotide V2 5′ UTR sequence and the exon 1b / exon 2 boundary are shown beneath the gene structure; the region shared across Variants 1–3 is indicated. **b.** *C9ORF72* transcript abundance (DESeq2-normalized counts) in post-mortem cerebellum from neurologically normal controls (n = 8) and *C9ORF72* repeat-expansion ALS (C9-ALS, n = 8), taken from GSE67196. Transcript levels are significantly lower in the C9ORF72 HRE samples (p-value < 0.001, two-sided Mann-Whitney U test). **c.** Aggregate ribosome P-site occupancy along the V2 5′UTR from >400 ribosome-profiling datasets (https://ribocrypt.org/). **Supplementary Figure S1** shows examples of individual experiments. Ribosome-protected fragments accumulate at the start-stop element (A74), the upstream translon AUG (uTranslon, A88) and the main translon start codon (A96); the A74 peak exceeds that at the main AUG. The grey region on the left indicates a GC-rich region. **d.** Secondary structure of the V2 5′UTR determined by SHAPE chemical probing. Left: experimentally-resolved secondary-structure model with nucleotides colored by SHAPE reactivity in the presence of 6 mM 1M7 chemical probe. Three stem-loops are resolved: SL1 (positions 9–40), SL2 (50–62) and SL3 (63–91). Right: per-nucleotide SHAPE reactivity (bars, mean ± standard error of the mean) with pairing-probability arcs below. **Supplementary Figure S3** shows the secondary structure at different concentrations of the chemical probe. **e.** Dual-luciferase reporter assay quantifying translation of V2 5′UTR variants in which large regions of the leader were replaced with an unstructured 14-nucleotide repeat sequence (M14). Constructs (top to bottom): wild-type (mc0); exons 1b and 2 represent M14 and wild-type C9ORF72 sequence, respectively (mc1, 1bM14 + 2WT); the inverse (mc4); and both exons replaced with M14 (mc3). Left: positional schematic showing each substitution in black and the respective sequence. Right: change in luminescence relative to wild-type, expressed as log2(mutant / WT) Renilla luciferase signal normalised to Firefly luciferase. Bars show mean ± standard error of the mean (n = 3 biological replicates with 3 technical replicates each). Statistical analysis: two-sided Student’s *t*-test vs. wild-type. A74, start-stop element at position 74; A88, upstream uTranslon at position 88; A96, main coding-sequence start codon; S72, frame-0 stop codon at position 72; SL, stem-loop; WT, wild-type. *P < 0.05, **P < 0.01, ***P < 0.001

The intronic region between exons 1a and 1b contains a hexanucleotide repeat (GGGGCC), which is present in 6-25 copies in healthy individuals, but can expand to >1,000 copies (**Figure 1a**), constituting a leading genetic cause of Amyotrophic Lateral Sclerosis (ALS) or Frontotemporal Dementia (FTD)^10^. Both ALS and FTD are fatal neurodegenerative diseases without a known cure. The hexanucleotide repeat expansion (HRE) impairs correct splicing of the gene, causing retention of the respective intron and translation of the hexanucleotide through repeat-associated non-canonical (RAN) translation ^11,12^, leading to toxic gain-of-function effects of the repetitive RNA and dipeptide. Intron-retention and RAN translation is thought to affect Variants 1 and 3, but not Variant 2.

However, the HRE does lower transcription of Variant 2 due to CpG methylation of the respective DNA region which contains Variant 2’s promoter region, causing a loss-of-function effect that aggravates the disease pathology even further ^1,13^. Indeed, re-analysis of published data from post-mortem cerebellum tissues from neurologically normal controls (n = 8) and *C9ORF72* repeat-expansion ALS cases (n=9) shows a highly significant reduction in total C9ORF72 transcript levels in the disease case (p<0.001; **Figure 1b**)^14^. Congruently, *C9ORF72* depletion in cellular models leads to severe phenotypes, i.e. axon growth defects, cilia misformation, impaired autophagy, and increased glutamate receptors, all of which impair neuronal health ^7,15^. While recent advances using antisense-oligonucleotide (ASO) mediated editing have successfully repressed HRE translation and the pathological expression of dipeptides, they failed to restore Variant 2’s transcript levels, leaving the loss-of-function aspect of the model unaddressed ^16^.

When examining published, aggregate ribosome footprints along the 5’UTR of the mature transcript for C9ORF72 Variant 2, we observed the high occupancy at position 74 on a start-stop element, which consists of only an AUG start codon immediately followed by a UGA stop codon - which was even higher than on the main translon’s initiation site (A96) or another short uTranslon (A88)(**Figure 1c**). This observation was true also when examining individual ribosome footprinting datasets (**Supplementary Figure S1**). Note that uTranslons and main translons were formerly known as uORFs (upstream open reading frames) and coding regions, respectively ^17^. We have recently characterized start-stop elements in detail, demonstrating that, together with uTranslons, they represent important elements of translation repression ^18^. The start-stop in C9ORF72 also showed the highest evolutionary conservation of the entire 5’UTR (**Supplementary Figure S2**), further underscoring its potential significance.

Based on these observations, we carefully probed sequence and structure elements in Variant 2’s 5’UTR for their impact on C9ORF72 translation, and discovered a complex, multi-layered regulatory system. Central to this regulatory system are ‘buffered repressors’ which ensure low baseline C9ORF72 levels that are resistant to fluctuations in translation propensity - consistent with C9ORF72’s critical function in neurons.

## Results

### Chemical probing resolves three stem-loops in the Variant 2 5′ UTR

First, we used experimental and computational methods to determine the secondary structure of Variant 2’s 5’UTR. To do so, we probed the *in vitro* transcribed leader by selective 2′-hydroxyl acylation analysed by primer extension with mutational profiling (SHAPE-MaP)^19–21^. The resulting per-nucleotide SHAPE reactivity profile, combined with reactivity-constrained structure modeling and the corresponding base-pairing probabilities, defined three stem-loops along the 95 nucleotide Variant 2 5′ UTR which we labeled SL1 to SL3 (**Figure 1d**). SL1, SL2, and SL3 spanned nucleotides 9 to 40, 50 to 60 and 63 to 91, respectively. The three stem-loops together fold approximately three-quarters of the sequence into base-paired structures, with single-stranded reactivity confined to the inter-stem-loop linkers and the 3′ region immediately preceding the Variant 2 main translon (**Figure 1d**). The computationally predicted SL1, SL2 and SL3 boundaries in the wild-type sequence were broadly consistent with the SHAPE-derived stem-loop assignments and consistent across variable concentrations of the chemical probe (**Supplementary Figure S3a,b**).

We noticed that SL1 coincided with the GC-rich segment in exon 1b that is specific to Variant 2 and which produced very few ribosome footprints in the aggregate profile shown in **Figure 1c**. As GC-rich regions are prone to reduced read coverage, in particular in short-read sequencing ^22,23^, we concluded that ribosome footprinting alone was insufficient to determine if ribosomes are bound to this region or not. SL2 spanned the exon 1b-2 boundary after nucleotide position 51 and represented the least structured element. SL3 was positioned to exon 2 which is common to the three major C9ORF72 transcript variants and much more conserved across vertebrates than exon 1b. SL3 enclosed the start-stop element (A74) and the AUG start codon of the uTranslon (A88) in structurally distinct positions: A74 sat in the unpaired loop/bulge, whereas A88 was in the paired stem.

### Unexpected changes in Variant 2 translation suggest complex regulatory relationships

Based on the observed secondary structures and ribosome occupancy at the start-stop and uTranslon (**Figure 1c, d**), we hypothesized that translation of C9ORF72 may be affected: the secondary structures or initiation at the start-stop or uTranslon may delay movement of the preinitiation complex (PIC) along the 5’UTR and therefore lower the rate of initiation at the main translon’s AUG. To test this hypothesis, we conducted a first broad test changing large parts of the 5’UTR. We cloned the Variant 2 5’UTR upstream of the renilla luciferase start codon into a single plasmid dual-luciferase reporter psiCHECK-2 such that the renilla translation reports 5’ UTR activity, with firefly luciferase expressed from an independent promoter on the same plasmid. Therefore, the luciferase signal we report is the ratio of renilla/firefly luminescence, controlled for transfection efficiency and gene expression levels. We modified either exon 1b, exon 2, or both into a sequence without predicted translation initiation sites or predicted basepairing (**Supplementary Figure S3c**). The position of the sequence changes and the resulting change in luciferase signal compared to the wild-type signal (and normalized to the internal firefly control) are shown in **Figure 1e**. Note that the figure shows log base 2 transformed data, i.e. an increase in translation (and luciferase signal) results in a positive, a decrease in a negative signal. The figure also shows the sequence of the reporter construct with black marks above indicating the region different from wild-type. **Supplementary Data S1** shows the raw data and corresponding RNA measurements.

Our natural expectation was that elements in both exon 1b (i.e. SL1, SL2) and exon 2 (i.e. SL2, SL3, the start-stop, and the uTranslon) may be translation inhibitory and their elimination would increase reporter translation. Indeed, substitution of exon 1b or 2 increased reporter translation ∼3- and ∼23-fold, respectively (mc1, mc4; **Figure 1e**; **Supplementary Data S1**), confirming that the elements in both exon 1b and, especially, exon 2 repress C9ORF72 translation. Surprisingly, however, when we replaced the entire 5’UTR with the unstructured/no-initiation sequence (mc3), the observed increase in luciferase signal was smaller than that observed for the substituted exon 2 (mc4), i.e. ∼16- vs. ∼23-fold. This observation was not due to changes in transcript levels (**Supplementary Figure S4**). While the observation may have been due to technical artifacts caused by the long unstructured RNA sequence, it also suggested a potential functional interaction between the elements in exon 1b and 2, i.e. the existence of a more complex regulatory network.

### Systematic mutant screening reveals several regulatory layers

To test if C9ORF72 translation may indeed be controlled by a more complex regulatory network, we first examined Variant 2’s 5’UTR carefully with respect to additional potential translation regulatory elements (**Figure 2a**). First, we mapped the three stem-loops (SL1 to SL3) discussed above with their position estimated from the SHAPE and structure prediction. Next, we split the annotation into the three reading frames, with frame 0 corresponding to the frame of the C9ORF72 translon and mapped the AUG start codons as well as stop codons into the respective frames (**Figure 2a**). The start-stop (A74) and the uTranslon (A88) were positioned to frames 2 and 1, respectively. The stop codon of the uTranslon overlapped with the AUG start of the main translon at position 96. A third stop codon in frame 0 (S72) overlapped with the AUG of the start-stop (A74).

**Figure 2.**
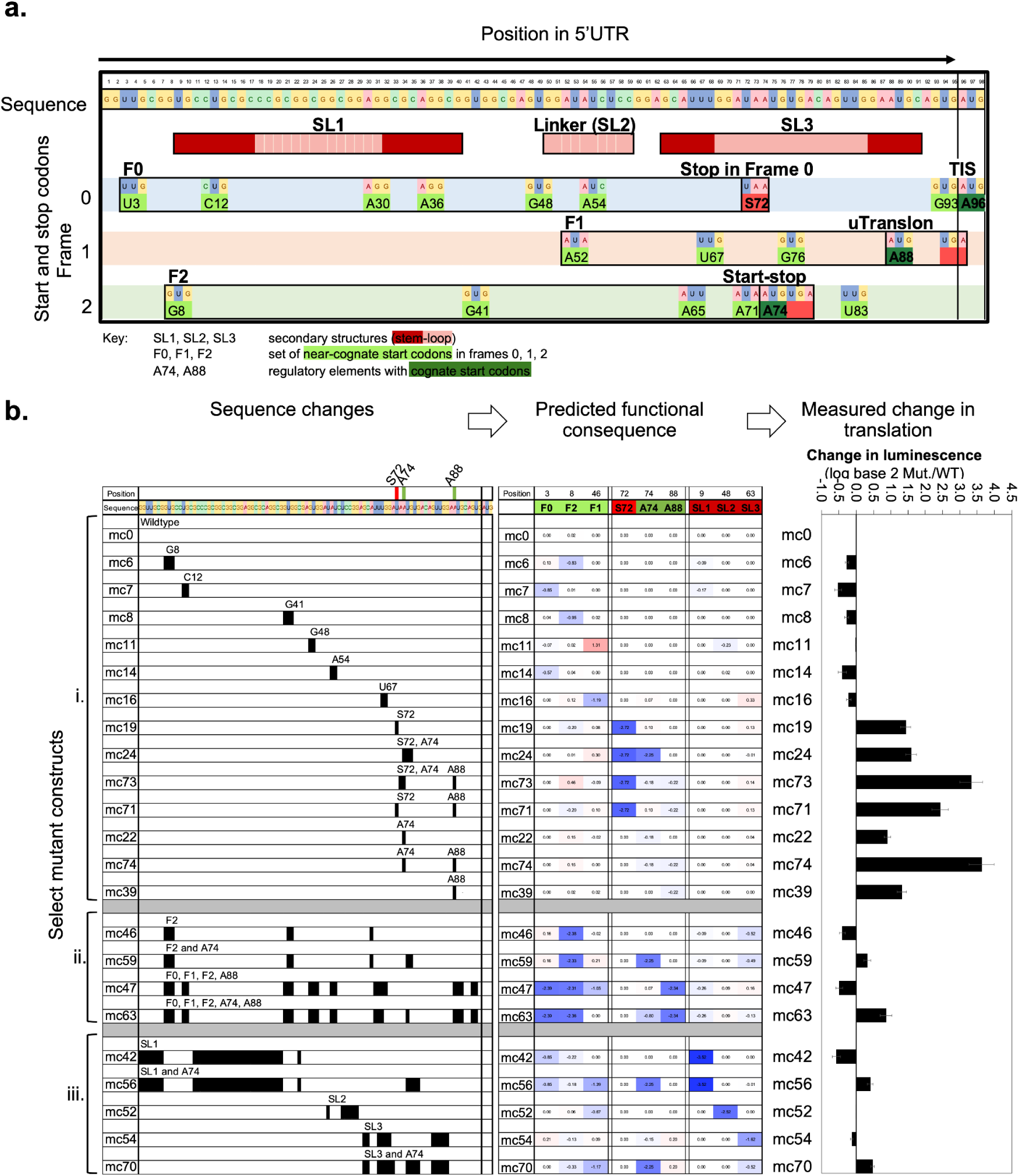
Systematic screening of the Variant 2’s 5’UTR suggests complex regulatory relationships. **a.** Predicted structure and sequence elements and their position in the 5’UTR of *C9ORF72* Variant 2. Magenta regions indicate predicted stem-loops (SL1, SL2, SL3). The predicted start and stop codons are sorted into the three frames (F0, F1, F2). Canonical (dark green) and near-cognate (light green) start codons were labeled with respect to their first nucleotide and position in the sequence, e.g. A74 denotes the A in the AUG of the start-stop element. **Supplementary Data S1** provides details on the strength of the initiation sites and predicted structures. **b.** Selected mutant constructs (n=23, total n=76) with modifications (black, left), consequence of the mutation on sequence/structure elements (middle), and resulting change in reporter signal (right). Mutant constructs were sorted into three groups: **i.** small sequence changes (<6 nucleotides) targeting individual start codons; **ii.** sequence changes targeting F0 to F2 in combination with the start-stop (A74); **iii.** Sequence changes targeting SL1 to SL3. Within each group, mutants were sorted from 5’ to 3’ with respect to the mutation’s position. For visualization, if a single unchanged nucleotide was surrounded by changed nucleotides, it would still be colored in black. The three major elements (S72, A74, A88) are indicated in the sequence. The middle panel shows the functional effect of the modifications on the elements. For visualization, the data was column Z-score normalized. Shades of red indicate an increase in value, shades of blue a decrease in value. A negative value (blue) for S72 indicates loss/mutation of the stop codon. A positive (red) value for F0, F1, F2, A74, and A88 indicates an increase in the strength of the translation initiation site(s) compared to wild-type; a negative value indicates the opposite. A positive value for SL1 to SL3 indicates that the predicted absolute free energy of the structure increases in the mutant construct, i.e. the structure becomes more stable; a negative value indicates the opposite. The panel on the right shows the change in reporter translation compared to the wild-type sequence (mc0)(log base 2 transformation), reported as mean and standard error of the mean from n=3 replicates. Details of all mutations and resulting reporter signals are provided in **Supplementary Data S1. Supplementary Figure S4** shows the RNA changes for select mutant constructs. A74 - start-stop element at position 74; A88 - uTranslon in frame 1; F0, F1, F2 - collection of near-cognate start codons in frames 0, 1, and 2; mc - mutant construct; Mut. - mutant; SL1, SL2, SL3 - Stem-loops; S72 - STOP in frame 0 at position 72; WT - wildtype

Finally, we identified 16 near-cognate start codons across all three reading frames of the wild-type 5’UTR with varying predicted strength of the translation initiation site (Kozak score). GUG at position 48 (G48) showed some ribosome coverage above background, indicating that ribosomes might indeed initiate at the site (**Figure 1c**). Most of the near-cognate start codons were located upstream of the stop codon in the respective frame, extending the set of possible initiation sites beyond that of the start-stop and uTranslon (**Figure 2a**). Below, we refer to the set of near-cognate start-codons in frames 0, 1, and 2 and upstream of the respective stop codon as F0, F1, and F2, respectively. F0, F1, and F2 have their respective stop codons in positions 72, 97, and 77, respectively (S72, S97, S77). Initiation at F0, F1, or F2 could also impact translation of the main translon, as reinitiation after their respective stop codon is unlikely due to the short distance between the stop codons and the main AUG. There were no predicted IRESes or RNA modification sites in the C9ORF72 Variant 2’s 5’UTR.

Next, we systematically tested the effect of these predicted elements on the translation of the main translon devising 76 constructs of the dual-luciferase reporter system described above. We introduced mutations targeting the near-cognate start codons across the three reading frames, the stop codon in frame 0 (S72), the start-stop element (A74), the upstream AUG-initiated uTranslon (A88), and the stem-loops (SL1, SL2, SL3). Mutations changed either single elements or combinations of elements. For each mutant construct, we reported the log base 2 transformed change in reporter translation compared to the wild-type sequence. **Supplementary Data S1/S2** list the precise mutant sequences, translation measurements and corresponding RNA data. RNA levels did not change substantially for the tested constructs (**Supplementary Data S1, Supplementary Figure S4**). Therefore, the observed change in luciferase signal largely corresponded to a change in translation.

**Figure 2b** shows a representative subset of the mutant sequences, organized into three groups: i. small changes targeting only one or two elements; ii. larger sequence changes targeting all near-cognate start codons in F0, F1, and F2, alone or in combination with A74 and/or A88; iii. larger sequence changes targeting SL1 to SL3 alone or in combination with A74. **Supplementary Data S2** shows the entire set of results for all 76 constructs.

While we attempted to design all mutations to be as specific as possible and to avoid secondary changes, doing so comprehensively was impossible as many elements overlapped, e.g. a change in a start-codon may also affect the secondary structural element position to the same region. Therefore, we developed a system in which we estimated the functional impact of a substitution not only on the element itself, but also on all other elements in the entire 5’UTR. To do so, we used the following proxies: for all canonical and near-cognate start codons, we monitored the change in predicted translation initiation strength (‘Kozak context’) and the start codon type; for all secondary structures (SL1 to SL3), we monitored the change in predicted free energy of the structure; and for the stop codon(s), we used a singular weighted score to denote the presence or absence of a stop codon. This approach also allowed us to distinguish between different extents of sequence changes, i.e. between completely abolishing an AUG by changing it into CCC versus just changing it into a near-cognate start-codon, such as CUG. All changes in functionality were recorded as the log base 2 ratio of the mutant versus wildtype element’s numerical value. Finally, to reduce the number of variables, we summarized the log base 2 ratios for all near-cognate start-codons into a single value F0, F1, and F2. The position of the sequence changes, the predicted quantitative functional impact of the change, and the resulting change in reporter translation are shown for select constructs in **Figure 2b**. **Supplementary Data S2** shows the entire dataset.

While far from perfect, this system allowed us to monitor the impact of any sequence changes onto all elements - simultaneously and in a quantitative way. One limitation of the system was that the functional impact of the stem-loops was captured only by the predicted change in free energy, which - in particular for very transient structures such as SL2 - was a poor reflection of its role in the sequence. Further, the Kozak score alone likely did not capture the full effect of start-stop disruption, as start-stops are distinct from uTranslons in their initiation properties ^18^.

Indeed, using this approach helped identify confounding secondary effects which in turn helped with the interpretation of the results (**Figure 2b, Supplementary Data S2**). For example, the changes in the mc11 construct (group **i**) eliminated the GUG in frame 0 at position 48 (G48), but also created a new AGG start codon in frame 1 at position 49 with a strong Kozak context, increasing the total initiation score for F1 while decreasing the score for F0. Further, two nucleotide changes in mc16 targeting the UUG in position 67 not only abolished the start codon, but stabilized SL3 (**Figure 2b, group i**). It also slightly increased the score for F2 due to improvements in the Kozak context for one of its start codons. In general, it was impossible to separate the effects of individual mutations in the SL3 region clearly with respect to affecting individual elements, as SL3 overlapped with the start-stop and uTranslon. While we focussed on the interpretation of changes in the start-stop and uTranslon, it cannot be ruled out that any observed effects arose from changes in SL3 instead.

Next, we examined the results to begin to understand how the regulatory elements influence translation (**Figure 2b**, **Supplementary Data S2**). We started with the simplest case, i.e. mutation of individual elements which resulted in no or little secondary changes (**group i**). We made several observations. First, most mutations of near-cognate start codons upstream of the stop codon in position 72 lowered reporter translation slightly, suggesting that the elements had a positive impact on translation of the main translon (**group i**: mc6, mc7, mc8, mc11, mc14, mc16). None of the mutations, except for the one in mc7, affected SL1’s predicted free energy which is localized to the same region. For example, the luciferase signal was *lower* compared to that of the wild-type when mutating the three near-cognate start-codons in frame 0 (C12, G48, and A54; **group i**: mc7, mc11, mc14). However, contrasting this observation, mutating the corresponding stop codon in frame 0, S72, resulted in an increase in reporter signal (**group i**: mc19). This increase in reporter signal upon mutation of the stop codon in frame 0 suggested that there was indeed initiation at the start codons in F0 and without S72, they would translate into an N-terminal peptide fused to the luciferase.

In comparison, mutations at or downstream of the stop codon S72, i.e. those of S72, A74, and/or A88, all increased reporter translation (**group i**: mc19, mc24, mc73, mc71, mc22, mc74, mc39). Note that except for mc24 (**group i**) where the start-stop’s AUG was changed into CCC, the other changes to A74 or A88 resulted in much smaller decreases in initiation potential of the respective start site, as the AUG start codon was changed to near-cognate start codons instead of abolishing it completely (**group i**: mc73, mc71, mc22, mc74, mc39). Nevertheless, some of these comparatively small changes, e.g. the individual mutations of A74 and A88 in mc22 and mc39, respectively, resulted in marked increases in reporter translation, supporting the expectation that both the start-stop and uTranslon were repressing C9ORF72 translation. Even more so, mutating both A74 and A88 to near-cognate GUG in mc74 had a de-repressive effect on reporter translation beyond what would have been expected by adding the individual signals.

Next, we hypothesized that the net positive effect of the near-cognate start codons upstream of S72 may result from them repressing the repressive role of A74 and A88: ribosomes initiating at upstream near-cognate start codons would be engaged in elongation and unable to initiate at the start-stop or at the uTranslon which therefore renders them not functional. We searched for support of this hypothesis by examining more complex mutant constructs where we mutated the upstream elements without and with simultaneous mutation of A74 or A88 (**group ii/iii, Figure 2b**). Mutating F2 alone decreased reporter translation (**group ii**: mc46), while the additional mutation of A74 led to an increase in reporter translation (**group ii**: mc59). This switch from a negative to a positive signal was even more extreme when mutating F0, F1, F2, and A88 without and with simultaneous mutation of A74 (**group ii**: mc47 vs. mc63). This observation could suggest that initiation at F0, F1, F2 may not repress A88 as much as it represses A74. Results for additional scenarios are provided in **Supplementary Data S2**.

Finally, we showed that SL1 and SL3 seemed to have an effect similar to that of the F0, F1, and F2: mutating SL1 or SL3 on their own decreased reporter translation, suggesting a net positive contribution in the wildtype context (**Figure 2b, group iii:** mc42 and mc54). Combining these mutations with mutation of A74 reversed the effect and increased reporter translation relative to the wildtype(**Figure 2b, group iii:** mc42 vs. mc56 and mc54 vs. mc70). Mutating SL2 on its own had no effect (**group iii**: mc52). However, despite carefully choosing the sequence substitutions eliminating the stem-loops, mutating the stem-loops still also affected the Kozak strength in F0, F1, and F2, which may have also caused the apparent switch in signal.

In sum, the results from the mutant screen confirmed the repressive roles of the start-stop (A74) and uTranslon (A88). F0, F1, F2, as well as SL1/SL3 suggested a net positive effect on translation when mutated on their own; however, combining their mutation with the mutation of A74 (and A88) suggested that this effect was net result of the structures and/or near-cognate start codons buffering the negative role of the start-stop and uTranslon. In other words, our results suggested extensive functional interactions between the elements in the 5’UTR.

### Quantitative modeling supports a complex regulatory system

Next, we devised a data-driven workflow to analyze all data in a quantitative, unbiased way and test if indeed the data supported functional interactions between different regulatory elements. To do so, we devised a modeling framework to estimate the effect of individual elements on translation of the main translon and select interactions between elements that may improve the modeling (prediction) outcome even further (**Figure 3a**).

**Figure 3.**
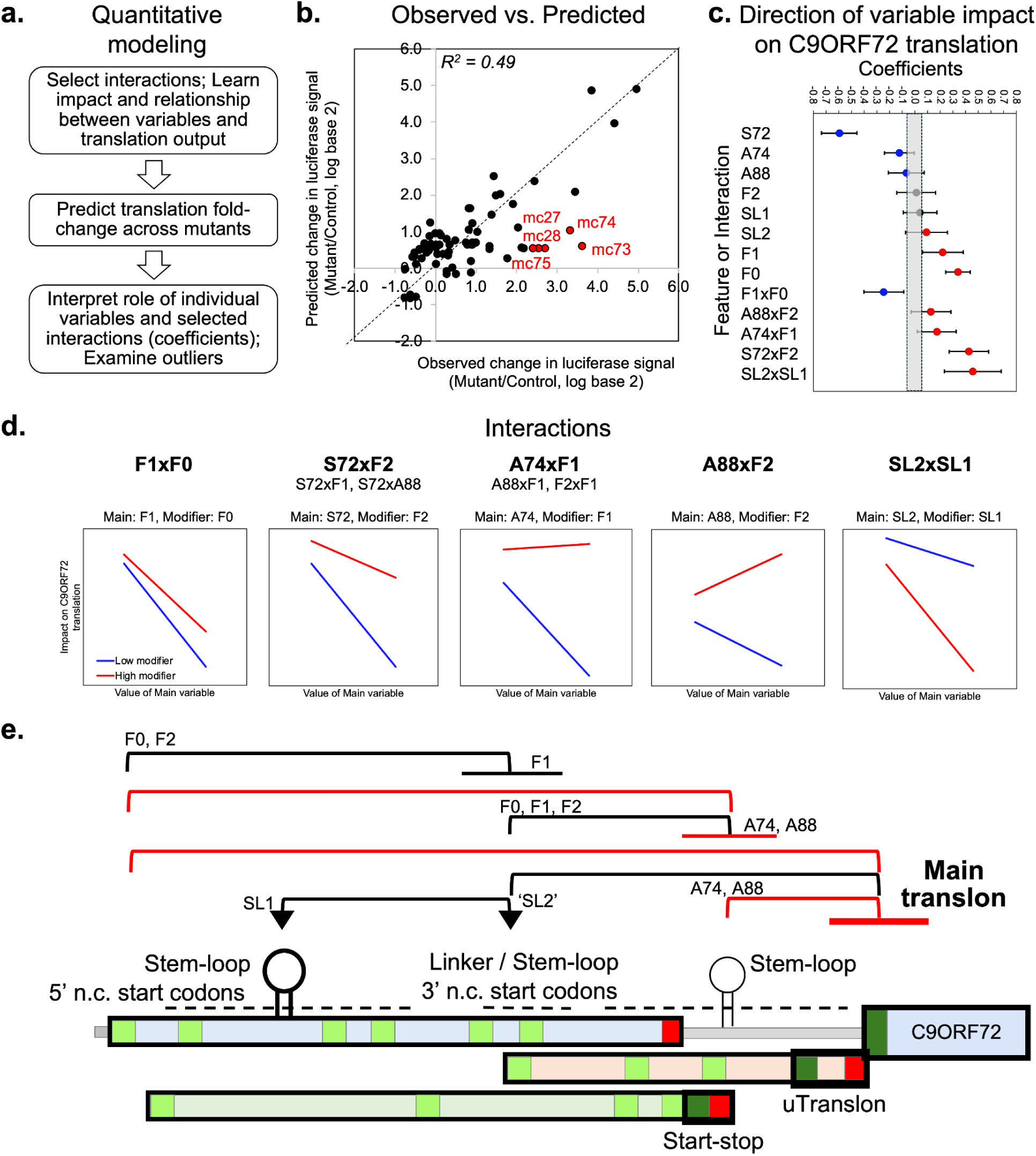
Quantitative modeling supports a multi-layered regulatory model governing C9ORF72 translation. **a.** Flowchart of modeling workflow. **b.** Relationship between observed and predicted changes in translation for all mutant constructs. Red indicates the mutant constructs with the largest residuals. Mc73, mc74, mc75 are further discussed in Figure 5. **c.** Coefficients for 8 individual variables and the 5 selected interactions, shown as mean and standard deviations from 100 bootstrappings. Blue indicates negative, red indicates a positive impact on C9ORF72 translation. The greyed out region indicates near-zero coefficients. **d.** Split-plot analysis illustrating one possible interpretation of the interaction. For each interaction, we selected a main and a modifier variable based on possible molecular mechanisms. We then estimated the impact (slope) of the main variable on reporter translation at high and low modifier values, where high and low were determined by the median modifier values. **e.** Putative model highlighting different regulatory layers inhibiting C9ORF72 translation. The model was derived from what is known about the effect of uTranslons and start-stops and from the modeling results. Top: inhibitory relationships indicated as source and target. Horizontal lines indicate inhibition; triangles indicate enhancement. Red lines indicate the specific feedforward loop tested in Figure 4. Bottom: graphic representation of the major regulatory elements. Further details are provided in **Supplementary Data S2.** A74 - start-stop element at position 74; A88 - uTranslon in frame 1; F0, F1, F2 - near-cognate start-codons in frames 0, 1, and 2; mc - mutant construct; R2 - Pearson’s correlation coefficient; SL1, SL2, SL3 - Stem-loops; S72 - STOP in frame 0 at position 72; u9 - nine-nucleotide uTranslon; WT - wild-type

We chose multiple linear regression for the ease of its interpretation, fully aware of its limitations, e.g. with respect to assuming linear relationships and independence between variables. Further, even with 76 constructs in the screen, the system was largely underdetermined which led us to drop the results for element changes that were uninformative, e.g. those for U83 and A96, or that highly correlated with the results for other elements, e.g. SL3. See **Methods** for details. Due to these limitations, we considered the outcome of the modeling exercise as a way to support or reject our manual interpretation of the data, rather than being the ultimate proof of existence for any regulatory model. The model does, however, provide a way to isolate the impact of individual elements and their combinations from each other in a more systematic and quantitative manner than what we were able to do manually.

To predict translation of the main translon across the 76 constructs, we assembled the translation and functional data for the main variables (SL1, SL3, F0, F1, F2, S72, A74, A88) as described in the **Methods**. For each modeling round, we used the Pearson correlation squared (R^2^) between the observed and predicted change in reporter translation as a measure of success. We believe that any more sophisticated error estimation would have gone beyond what the size of the dataset allowed.

The method first modeled the impact of (i.e. coefficients for) all main variables, i.e. the stem-loops, near-cognate start codons, start-stop and uTranslon, and then test for model improvements (R^2^ between observed and predicted) when including interactions. The interactions were tested in the order of stability of their contribution (coefficient). The selection procedure excluded interactions with high correlation (redundancy) with main variables or interactions that were already part of the model. We then refined the final model manually with respect to easier interpretation as described in the **Methods**. The final selected interactions were: F1xF0, A88xF2, A74xF1, S72xF2, and SL2xSL1. While the selection of the variables was performed on the residuals of a primary model involving just the main variables (as should be), the final model discussed here results from simultaneous fitting of all main variables and selected interactions. We then used the model to estimate coefficients for each main variable (element) and five pairwise interactions.

**Figure 3b** shows the result for the 76 constructs with respect to observed and predicted translation change of the reporter (labeled mc0 to mc75). The final R^2^ between observed and predicted translation changes was 0.49, suggesting our model captured nearly half the observed changes in translation. Red font in **Figure 3b** labeled the five least-well predicted mutants (largest residuals), i.e. those for which the model did not capture the regulatory impact. Of these, we discuss mc74, mc75, and mc73 more in detail below.

**Figure 3c** shows the model’s coefficients for the main variables and the five selected interactions estimated in the final model. Positive coefficients, denoting a positive impact on translation, are colored in red, negative coefficients in blue, very small coefficients are greyed out. Note that the sign rather than the absolute value of the coefficient should be considered, as at least for the stop codon (S72) the actual value is entirely arbitrary.

First, we observed that the positive coefficients for F0, F1, and SL2 and the negative coefficients for S72, A74, and A88 confirmed our manual interpretation of the screening results (**Figure 2b**, **Figure 3c**): the near-cognate start codons in frame 0 and 1, as well as SL1 and SL2 had a net-positive (enhancing) effects on C9ORF72 translation, while the start-stop and the uTranslon had expected repressive effects. Again, the large negative value for S72’s coefficient might be due to the value we assigned for the mutation of a stop codon. The comparatively small negative values for the coefficients for A74 and A88 might be due to their role in several of the interactions.

Second, the model also recapitulated the positive effects of the mutations in F0 and F1 discussed above. Third, F2 and SL1 resulted in very small coefficients, likely due to their strong involvement in the interactions. Indeed, when modeling only the main terms without interactions, SL1 had the expected non-zero coefficient (**Supplementary Figure S5**). Finally, the model reported a small but positive effect of SL2 which we had not noticed in our manual interpretation of the data.

Next, we examined the five interaction terms with non-zero coefficients which the workflow had selected (**Figure 3c**). From a modeling perspective, the interactions are undirected, e.g. F1xF0 is the same as F0xF1. Further, two interactions, S72xF2 and A74xF1, showed high correlation with other interactions, as listed in **Figure 3c**. With the present data, we could not say which interaction the signal came from and what the direction (if any) of the interaction would be. Therefore, the interpretation was guided by our specific hypotheses. We used a split-plot analysis to help interpretation of specific interactions (**Figure 3d**). To do so, we defined one of the interaction partners as the main variable (main), whose impact on the main translon’s translation was modified by the second interaction partner (modifier). The impact of the main variable was estimated by the slope using simple linear regression. Note that the direction of the slope may differ from that obtained in the multiple linear regression models discussed above due to the presence of confounding variables. We then examined the impact of the main variable with respect to the changes in the slope at high or low values of the modifier, splitting the data into two sets.

For F1xF0, we hypothesized that initiation at the F0 start codons may lower initiation at the F1 start codons, as the former were located upstream of the latter (**Figure 2a**). Indeed, F1 was negatively correlated with reporter translation for both high and low F0 values (F1xF0; **Figure 3d**), however, the impact (slope) was less negative at high initiation at F0 (red line, F1xF0; **Figure 3d**). Similarly, the stop codon in frame 0 (S72) always had a negative impact on reporter translation, but less so when initiation at F2 was high (red line, S72xF2; **Figure 3d**). Note that due to intercorrelation between interaction terms, the same argument holds true for high values of initiation at F1. These results quantitatively supported the notion that initiation at the upstream near-cognate start codons reduced initiation at the other start codons, i.e. the near-cognate start codons were reducing each other’s impact.

Further, the interactions A74xF1 and A88xF2 (**Figure 3d**) strongly supported the notion that initiation at the near-cognate start codons reduced initiation at the start-stop and uTranslon, therefore lowering their repressive effect on the main translon. High values for the modifiers F1 and F2 substantially reduced the negative slope (impact) between reporter translation and A74 and A88, respectively. In fact, they even flipped the sign and rendered the impact (slope) of A74 and A88 on reporter translation slightly positive. Due to intercorrelation between the two interaction terms and other interactions, the signal could also come from F1 impacting A88, or F1 and F2 impacting each other (**Figure 3d**). While impossible to deconvolute in every detail, the results are consistent with a model in which initiation at the near-cognate start codons negatively affect initiation at other start codons, including the start-stop and the uTranslon.

Finally, we examined the one selected interaction between the stem-loops, SL1 and SL2 (SL2xSL1; **Figure 3d**). Note again that SL1 was a very stable structure, while SL2 not as much. Therefore, the measure of a free energy change might very insufficiently reflect the change in SL2 function. Further, we do not know of a specific molecular mechanism by which one stem-loop might influence the other, therefore, we simply tested the pragmatic interpretation of the upstream stem-loop (SL1) modifying the impact of the downstream stem-loop (SL2). However, the relationship could also be the other way around or undirected. We found that the overall negative impact of SL2 on C9ORF72 translation was enhanced by a strong SL1; a weak SL1 stem-loop also led to weak correlation between SL2 values and C9ORF72 translation. In other words, SL1 and SL2 appeared to positively affect each other’s function.

We assembled these results into a graphic illustration of the translation regulation of C9ORF72 Variant 2 (**Figure 3e**). In the model, the near-cognate start codons, start-stop and uTranslon all repressed C9ORF72 translation, providing one possible reason for the very low baseline expression levels observed for the gene in brain cells ^9^. However, the elements also impacted each other, in particular the upstream near-cognate start codons reduced, i.e. buffered, the effect of the downstream canonical inhibitory elements, namely the start-stop and uTranslon. The stem-loops further modified the model by interacting with each other and possibly with the uTranslons. These regulatory relationships formed several feedforward loops, specifically negative or incoherent type 2 feedforward loops: two elements both negatively affect the target (C9ORF72 translation), but one element also negatively affects the other, effectively buffering its impact. One such example involves F2 and A74 and is highlighted in red in **Figure 3e**.

### A chimeric reporter probes one of the feedforward loops

The quantitative modeling supported what we had hypothesized in the manual interpretation, i.e. that the observed net positive effect of the mutations of F0, F1, and F2 may be due to their interaction with A74 (or A88), namely repression of the start-stop’s (or uTranslon’s) negative effect on the main translon. On their own, F0, F1, and F2 repress reporter translation, forming incoherent feedforward loops involving a direct and an indirect pathway.

To test this interpretation, we devised a way to formally probe the existence and functionality of such incoherent feedforward loops in relation to F2 and A74 (**Figure 3c, 4a**). Specifically, we expected both F2 and A74 to repress the main translon directly, but F2 would also reduce (buffer) the A74’s function, forming an indirect pathway to the main translon (F2->A74->main translon) which was incoherent with the direct pathway (F2->main translon), through a net increase in translation (less repression).

To probe this model in a C9ORF72-like system but with minimized confounding effects, we devised a chimeric reporter construct in which we kept the original sequence for C9ORF72 exon 1b, but substituted the sequence for exon 2 with the 3’end of the ACTB 5’UTR which is thought to have no notable translation regulatory elements (**Figure 4b**). Further, we carefully changed all near-cognate start codons in the exon 1b part of the construct, leaving SL1 as intact as possible. **Supplementary Figure S8** shows this ‘Base’ construct and its by-frame properties: F0, F1, and F2 as well as any elements in the exon 2 part of the C9ORF72 5’UTR were eliminated, but SL1 and other exon 1b features were still present. At position 74, the equivalent to A74 in the wild-type C9ORF72 sequence, the Base construct has a cytosine, i.e. it is C74.

**Figure 4.**
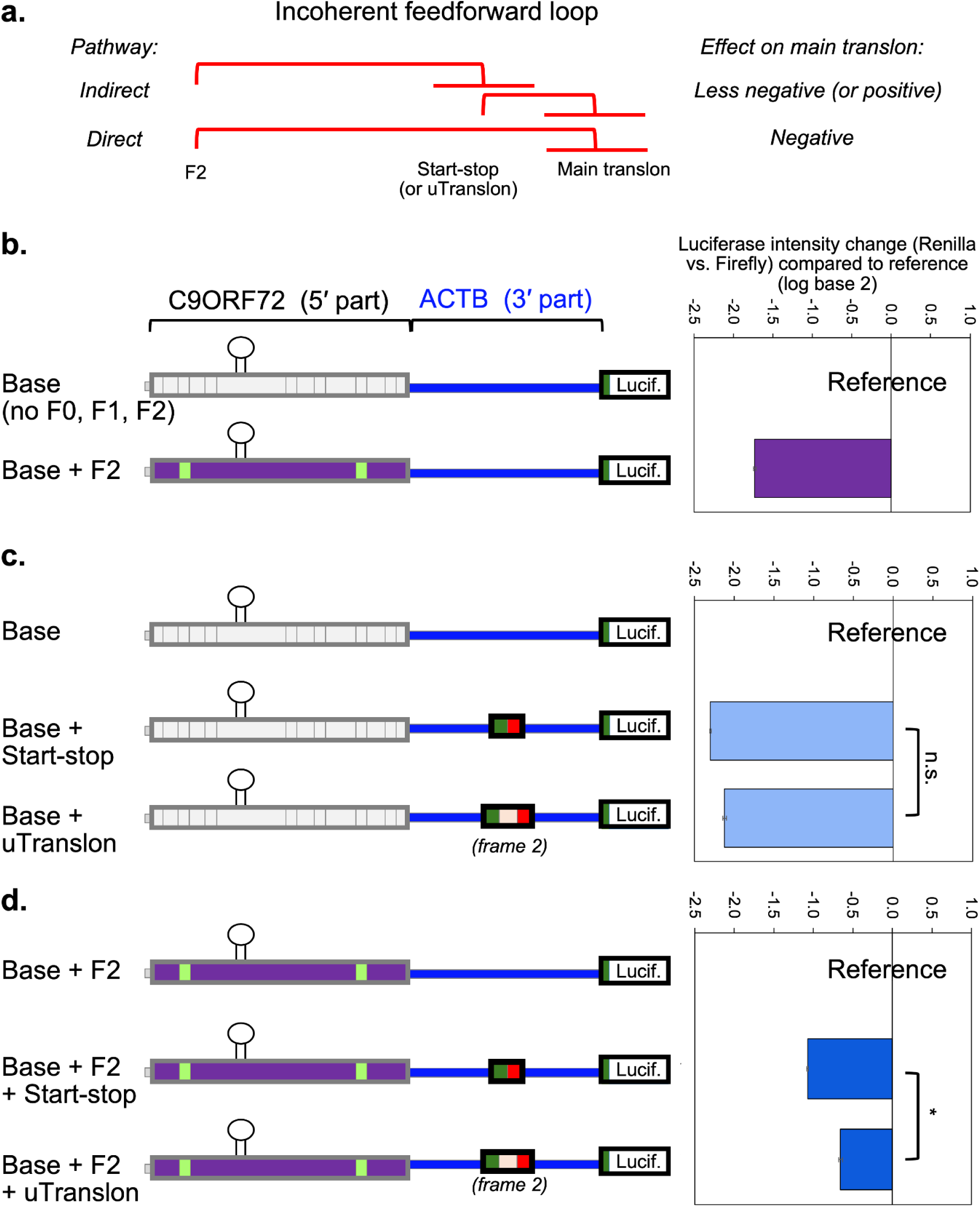
A chimeric reporter probes one of the feedforward loops rendering a buffered repressor system. **a.** Schematic of feedforward loop isolated from proposed regulatory model, involving F2 and the start-stop/uTranslon. **b.-d.** Chimeric reporter probing the buffering function of the feedforward loop illustrated in **a.**. The chimera comprises the first 51 nucleotides of C9ORF72 V2 5’UTR without any F0, F1, or F2 start codons, fused to the last 44 nucleotides β-actin (ACTB) 5’UTR driving the Renilla luciferase reporter (‘base’). The graphs show changes in reporter expression of the respective constructs relative to the reference construct. In panels **b.** and **c.** the reference is the ‘base’ chimera; in panel **d.** The reference is the ‘base’ chimera with the F2 start codons added back in. Panels **b.** and **c.** show the isolated repressive effects of F2 and the start-stop (or 9 nucleotide uTranslon) on the main translon, respectively. Panel **d.** shows the effect of the start-stop (or 9 nucleotide uTranslon) on the main translon, *in the presence of* F2. Translation repression is significantly reduced in panel **d.** compared to panel **c**. (p-value < 0.001, paired t-test).Error bars represent the standard error of the mean from n=6 replicates. F0, F1, F2 - near-cognate start-codons in frames 0, 1, and 2; Lucif. - luciferase; * - p-value < 0.05

We now systematically re-introduced elements to probe for their individual and combined effects as expected in an incoherent feedforward loop. Note that we focussed on re-introducing F2 and not F0, as F0’s regulatory scenario was much more complex: frame 0’s stop codon (S72) was located to the exon 2 region and overlapping with the start-stop’s start (A74), hence the presence/absence of the stop codon would not only impact F0’s function but also the start-stop.

Indeed, re-introducing F2 into the Base construct reduced reporter translation compared to Base (**Figure 4b**), setting a value for the direct pathway. The direct effect of F2 on the main translon was repressive, as expected. Further, the direct effect of a start-stop or a 9-nucleotide uTranslon re-introduced at position 74 into the Base construct also repressed translation of the main translon (**Figure 4c**) confirming the elements’ negative impact on translation: both the start-stop and the uTranslon reduced reporter translation about 4-fold (**Figure 4c**). Note that this engineered uTranslon introduced at position 74 is different from the uTranslon at position 88 in the wild-type C9ORF72 5’UTR.

To examine how the impact of the start-stop/uTranslon changes depending on modifier presence, we considered a new reference construct, i.e. the Base construct with F2 included (Base+F2; **Figure 4d**). When introducing either the start-stop or the uTranslon into the Base+F2 construct, we observed a buffering effect: introducing the start-stop or uTranslon reduced translation only 2-fold or less, i.e. substantially less than what we had observed in **Figure 4c**. If considering absolute translation changes instead of fold-changes, the buffering effect was even stronger. The results were not due to changes in RNA levels (**Supplementary Data S1**). These results support the notion of a functional, incoherent (type 2) feedforward loop participating in translation regulation of C9ORF72 translation: upstream F2 attenuates the repression by the downstream start-stop.

### Specific single-nucleotide changes cause unexpected effects

Finally, we examined more closely the set of constructs marked in red in **Figure 3b**. These constructs provided translation increases larger than those predicted by the final model, i.e. their translation regulatory network might involve additional components not captured yet. The outliers comprised two groups: constructs in which the start-stop had been extended to longer uTranslons (**Supplementary Data S2)** and single nucleotide changes affecting A88 with or without also changing A74 or S72 (mc73, mc74, mc75).

We examined the single-nucleotide changes more closely and compared them to similar sequences, zooming into the start-stop/uTranslon region (**Figure 5a**) as the remaining sequence was untouched in these constructs. We noticed that, for example, completely abolishing the uTranslon’s AUG start codon (mc40) had a smaller effect on reporter translation than changing it into a near-cognate start codon such as GUG (mc39). This difference suggests that loss of AUG initiation alone does not explain the response. One possible explanation for the effect lies in the fact that the sequence change in mc40 also destabilized SL3. Another explanation may be that the uTranslon’s AUG is essential for its repressive function and cannot be replaced by a near-cognate start codon. A similar observation applied to the start-stop (A74): changing its AUG to a GUG (mc22), which is a highly initiation-favorable near-cognate start codon has a similar effect to changing the AUG to a much less initiation-favorable ACC (mc31). Changing the AUG start codon at both the start-stop and the uTranslon to GUG (mc74) had an even stronger effect (∼10-fold change) which was larger than the sum of the individual effects which caused ∼2-fold changes (mc22, mc39). Structural probing of mc22 and mc39 using SHAPE did not reveal any notable differences to the SHAPE-constrained structure of the wild-type sequence (**Supplementary Figure S6**). This synergism suggested an interaction between the elements that we had not detected before.

**Figure 5.**
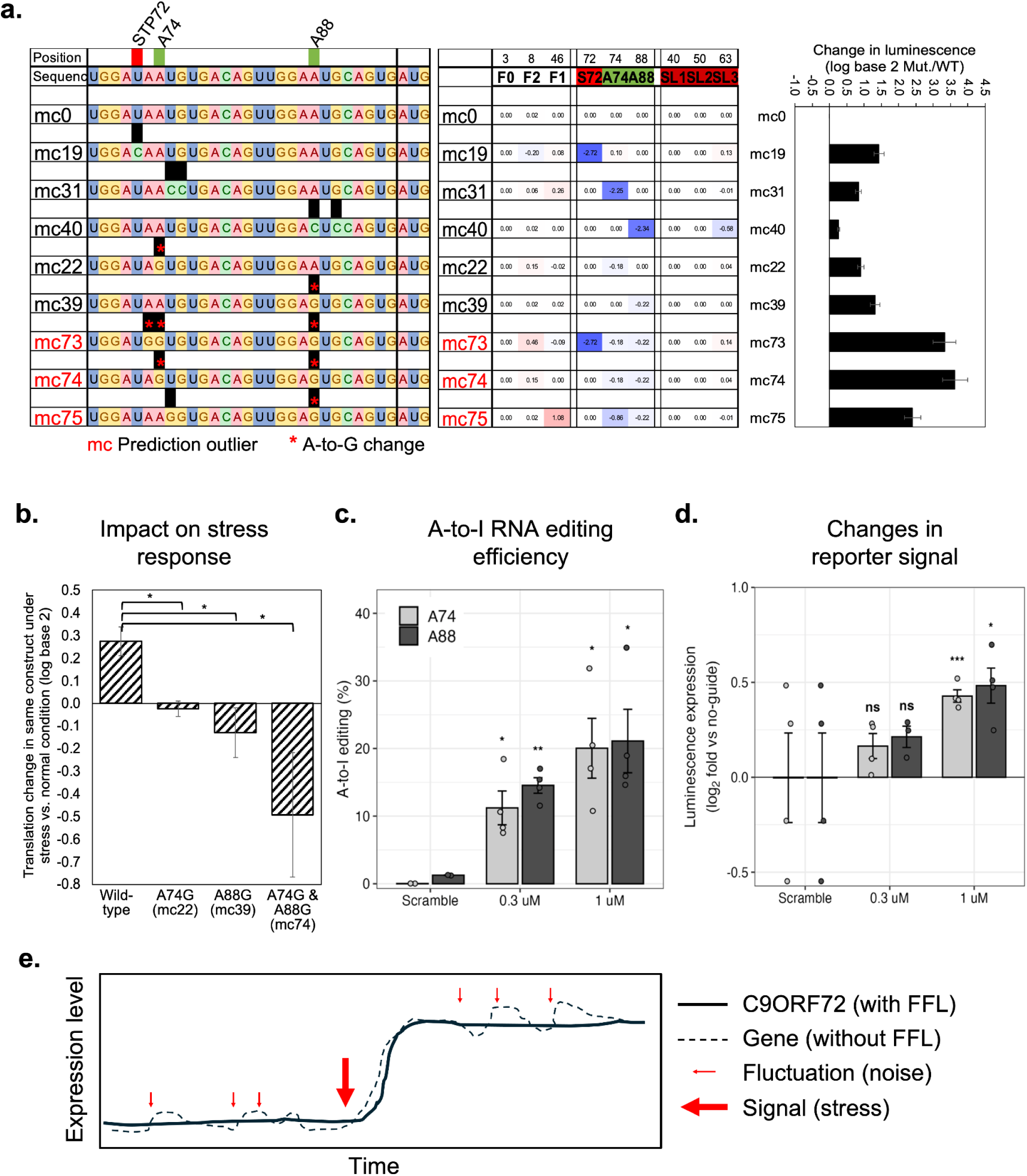
Select mutants affecting the start-stop and uTranslon. **a.** Selected mutant constructs with single-nucleotide changes. The three panels show the sequence changes (black), the functional consequence (blue and red) and resulting change in reporter translation (bar graph) similar to what is shown in Figure 2b. All selected constructs have changes only in the start-stop (A74) and uTranslon (A88) region. Construct names in red denote model outliers marked in Figure 3b. **Supplementary Figure S6** shows the SHAPE structure for mc22 and mc39. Error bars represent the standard error of the mean from n=3 replicates. **b.** Translation response to thapsigargin induced protein misfolding stress. The graphs show the log base 2 transformed changes in reporter construct translation upon 4 hour thapsigargin treatment for the wild-type and three mutant constructs. Error bars represent the standard error of the mean from n=3 replicates. **c.** Successful A-to-I editing at positions 74 and 88 in the C9ORF72 V2 construct. Editing at A74 and A88 were detected in HEK293T cells using targeted amplicon sequencing, with a scramble antisense oligonucleotide as control. Error bars represent the standard error of the mean from n=4 replicates. **Supplementary Figure S9** shows the successful positive editing control. **d.** A-to-I editing induced change in reporter translation. We measured dual-luciferase reporter translation (Renilla/Firefly, log₂ fold relative to no-guide) upon editing performed as in panel **c**. Error bars represent the standard error of the mean from n=4 replicates. **Supplementary Figure S10** shows the broader sequence context (specificity), the results for different ASO concentrations, and sequencing coverage for the respective editing sites. **e.** Graphic illustrating the effect of an incoherent feedforward loop (FFL) on gene expression. The FFL allows for gene expression levels to be adjusted in response to a signal, e.g. stress conditions under which PIC availability changes, but it is more robust to smaller signals, such as those coming from micro-fluctuations. A74 - start-stop element at position 74; A88 - uTranslon in frame 1; FFL - feedforward loop; mc - mutant construct; PIC - preinitiation complex; WT - wild-type; \**P* < 0.05; \*\**P* < 0.01; \*\*\**P* < 0.001; n.s., not significant.

To probe the role of the GUG mutations replacing the AUG start codons in the start-stop and uTranslon further, we examined the respective constructs under conditions of compromised translation initiation capacity, i.e. when the presence of an AUG is particularly important. To do so, we tested the constructs under protein misfolding stress induced by thapsigargin. The thapsigargin treatment triggers a response cascade which involves lowered translation initiation due to phosphorylation of the initiation factor eIF2α (**Supplementary Figure S8**). We observed that - in contrast to a global translation decrease - the wild-type C9ORF72 reporter was slightly induced under stress (∼1.2 fold; **Figure 5b**). This induction was completely abolished in the absence of the start-stop (mc22) or the uTranslon (mc39). Even further, reporter translation decreased when both the start-stop and uTranslon had been modified (mc74).

Having established that the upstream regulatory elements have the potential to repress translation of the main translon, we asked whether changing the corresponding nucleotides in the transcript would have the same effects as observed when mutating elements at the DNA-level in the reporter constructs. If successful, such an approach would provide the proof-of-concept for a clinical approach to increase C9ORF72 expression levels without permanently changing the genome and to alleviate the loss-of-function aspect observed in C9ORF72-linked ALS and FTD. Specifically, the DNA-level A-to-G substitutions discussed above would correspond to adenosine-to-inosine (A-to-I) RNA editing performed by endogenous ADAR enzymes. Inosine in turn is read as guanosine by the translation machinery.

Therefore, we designed an experiment to probe whether directed RNA editing at A74 or A88 of the unmodified reporter would relieve reporter repression similar to what we had observed for the mutant constructs (mc22, mc39, mc74; **Figure 5a**). First, we established the approach using a positive control construct in which the psiCHECK-2 ACTB 5’UTR reporter construct comprised a UAG stop codon immediately following the luciferase’s AUG start codon. We co-transfected HEK293T cells with the construct, an ADAR p150 expression plasmid, and a nuclease-resistant antisense oligonucleotide (ASO) targeting the UAG’s adenine. Successful editing converted the stop codon into a UGG and produced a luciferase signal (**Supplementary Figure S9**).

Next, we repeated the editing experiment using the wild-type psiCHECK-2 C9ORF72 5’UTR reporter and ASOs targeting A74 or A88. We used the corresponding scrambled ASO as the control and defined reporter expression in cells receiving p150 without an ASO as the baseline. Amplicon sequencing detected editing at A74 and A88 in cells treated with the A74- and A88-targeting ASO at ∼20% (**Figure 5c**). Editing remained at or near background levels in the scrambled ASO and no-ASO conditions (**Figure 5c**). Even more excitingly, these edits led to an up to ∼40% increase (1.4-fold) in luciferase signal (**Figure 5d**), supporting the notion that C9ORF72 translation increases in edited RNA similar to what we observed in mutant constructs. Because each matched comparison differed only in ASO sequence, these results support a sequence-dependent effect associated with the targeted editing rather than an effect of p150 expression alone. Editing efficiency increased with ASO concentration and was also highly specific, with little offtarget effects (**Supplementary Figure S10**). Therefore, the RNA editing mimics the A-to-G substitutions examined in mc22, mc39, and mc74 (**Figure 5a**).

## Discussion

We present an analysis demonstrating complex translation regulatory modes governing expression of *C9ORF72* Variant 2, which is the dominant transcript isoform at ∼70-100% of the gene’s transcripts in neurons ^8^ (**Figure 1b**). While the intronic hexanucleotide repeat expansion in C9ORF72 has gained much attention due to its detrimental gain-of-function effect in C9ORF72-linked diseases, the concurrent loss-of-function effect, i.e. lower C9ORF72 expression levels, has been much less studied. Even more so, very little is known about the regulation of the gene under normal conditions. In this work, we delineate a complex regulatory system that may keep C9ORF72 translation at low, but likely constant levels. Congruently, multiple studies have shown that the protein is required at a narrow physiological range and both over- and underexpression of the gene produces pathological cellular phenotypes ^24–28^. Low baseline translation is conceivable given C9ORF72’s role as a guanine exchange factor in endosomes/lysosomes which may be sensitive to high doses of the protein.

Further, we identified a multi-layered ‘buffered repressor’ system consisting of several incoherent feedforward loops which could form the basis for comparatively constant translation (**Figure 5e**). An incoherent feedforward loop is a regulatory circuit in which an input signal triggers both a direct and an indirect pathway, and the direct and the indirect pathway have opposite effects on the output ^29^. In the case of C9ORF72 translation, one incoherent feedforward loop is formed, for example, by F2 and the start-stop (**Figure 4a**). F2 has a direct inhibitory effect on the main translon: the more initiation occurs at the upstream near-cognate start codons, the fewer ribosomes initiate at the main translon. However, in the indirect pathway, initiation at F2 also reduces initiation at the start-stop which in turn lowers its repressive effect on the main translon compared to the start-stop’s effect when acting on their own. Therefore, the effect of the start-stop on the main translon is dampened or ‘buffered’.

Note that this buffering is different from simple redundancy seen in a system where multiple circuits regulate the same output. It is also different from translation-level buffering described in the literature ^30–33^. In those publications, buffering refers to the gene level, typically comparing transcript abundance with translation estimated from ribosome footprints: the studies report that variability in the transcript level of specific genes is often reduced (buffered) at the level of translation of the corresponding genes. Importantly, the buffering described here refers to different elements within one transcript: it describes the attenuation of one element’s repressive effect by another element.

Incoherent feedforward loops, in general, have the intrinsic property of allowing a response to an input signal without being as sensitive to noise or small fluctuations as other systems ^34^(**Figure 5e**). We speculate that this property might be particularly important in neuronal cells where translation capacity may fluctuate over time and across the cell ^35,36^, but C9ORF72 expression might still be required to remain constant. In other words, in addition to providing a fail-safe system with multiple backup circuits, C9ORF72’s translation regulatory system may also ensure expression levels that are as constant as possible in a highly variable cellular environment (**Figure 5e**).

To delineate C9ORF72’s translation regulatory system (**Figure 3e**), we first employed a mutant screen which measured translation changes via a reporter construct. We then developed a scheme to encode the sequence changes (mutations) as predicted quantitative consequences on the function of the respective elements. With this scheme, we were able to monitor the impact of a mutation on all sequence elements simultaneously, including cases in which a sequence change created new start codons in addition to destroying other elements. Finally, we used linear regression to correlate all measured translation changes with the main elements and select interactions within a single model. This workflow resulted in a model which could explain about half of the observed changes in C9ORF72 translation. While we designed this workflow to be as quantitative and data-driven as possible, it still faces limitations: mutant data was limited, variables may have been coupled, and the assumption of linear relationships may have not held. Despite these limitations, the framework is highly generalizable and can enable systematic analysis of the role of regulatory elements.

We then integrated the results of the automated workflow with our manual interpretations to derive the final regulatory model described in **Figure 3e**. In essence, the upstream near-cognate start codons, primarily in frames 0 and 2, but also in frame 1, suppressed translation of the main translon - as was expected - but also modified the downstream elements in their function, namely the start-stop at position 74 and the uTranslon at position 88. A stable secondary structure, i.e. a stem-loop in the Variant 2 specific region (SL1) further modified the system, e.g. by interacting with SL2. Our data did not support the notion that SL1 directly enhances translation, contrasting what had been proposed for other secondary structures ^37^. We found that SL1’s net enhancing effect was exerted through context-dependent, indirect action.

While our results clearly demonstrated SL2’s role in C9ORF72 regulation, it might not act through its structure, as the RNA secondary structure is fairly transient (**Figure 1d**). One possibility is that the SL2 region may act through providing a binding site for an RNA-binding protein. Indeed, HNRNPF, an RNA-binding protein with known function in neurodevelopment and neurodegeneration, is predicted to bind to the SL2 region (**Supplementary Table S1**)^38,39^. The potential HNRNPF effect on C9ORF72 translation is even more intriguing given that the hexanucleotide repeat expansion in diseased cells has been shown to sequester HNRNPF ^39^. However, comprehensive evaluation of the role of RNA-binding proteins in C9ORF72 regulation goes beyond the scope of this analysis.

The presence of and interactions between multiple translation regulatory elements, i.e. uTranslons, within one 5’UTR have been described for many genes, most notably perhaps ATF4 and GCN4 ^40–43^. In these cases, the arrangement of the uTranslons enables strong translation induction of the genes under conditions of reduced PIC availability, i.e. in response to cellular stress, when overall translation initiation shuts down. In the case of C9ORF72 the situation is different: the arrangement of the start-stop and uTranslon (as well as F0, F1, and F2) creates a regulatory model which leads to only little induction of C9ORF72 translation under stress, as shown in our data (**Figure 5b**) and by literature ^44^. Our data suggests that this behavior is still dependent on the start-stop and uTranslon (**Figure 5b**).

Finally, we exploited our findings to demonstrate the ability to alleviate the loss-of-function effect observed in C9ORF72 HRE cells. Specifically, we expanded upon the observation that changing a single nucleotide in the AUG in the start-stop and uTranslon to create a GUG substantially increased translation of the main translon (**Figure 5a**). We demonstrated that ASO-guided, ADAR-based RNA editing of the same sites can reproduce the effect observed for mutant constructs (**Figure 5c,d**)^45^. Such editing might occur naturally as a way to regulate the gene’s expression. However, screening published datasets, we found no evidence in support of this hypothesis (**Supplementary Figure S12**), indicating that our ASO-based editing approach installs a *de novo* edit rather than tuning a natural event. While the success of our RNA-editing is very promising (**Figure 5c,d**), several refinements, e.g. increased efficiency, modification of the endogenous gene, and cell-type specificity, would be necessary beyond this first proof of concept.

Our results provide a conceptual route towards counteracting C9ORF72 loss-of-function in the ALS/FTD mutant case by increasing translation of the main translon. Such RNA editing could complement recent work that addressed the gain-of-function arm of the disease by suppressing the production of HRE-derived dipeptide repeats through mutation of an upstream CUG start codon to CCG, while preserving repeat-containing RNA expression^16^. The approach is also conceptually similar to what has been shown recently for IRF6, where site-directed RNA editing of an inhibitory uTranslon relieved translation repression ^46^. Finally, instead of RNA-editing, similar effects on translation could in principle be achieved permanently by CRISPR-based editing of uTranslon elements in the corresponding DNA, an approach that has been used in plants to derepress downstream translation ^47^ and is now explored for its use in human disease ^48^.

In sum, to the best of our knowledge, our work reports for the first time a system of incoherent feedforward loops regulating the dominant transcript variant of C9ORF72, elucidated with the help of quantitative scoring to model the consequences of sequence changes. Further, it highlights an aspect of gene expression regulation that complements the large body of work investigating induction and repression of expression: regulation of a gene whose expression is required at low but fairly constant levels (**Figure 5e**). Finally, our work demonstrates that ADAR-based single nucleotide edits of native regulatory start codons can increase translation of a gene, similar to what has been shown for other examples ^46^ — a de-repression strategy that points to RNA editing as a general route to tune protein output.

## Materials and Methods

### Sequence and sequence analysis of the 5’ UTR of C9ORF72 Variant 2

Throughout the work, we refer to the C9ORF72 Variant 2 5’UTR (V2) as sequence NM_018325.5, ( ENST00000380003.8, C9ORF72-203) downloaded from Ensembl.org, comprising 98 nucleotides including the AUG start side of the main translon. We verified the correct transcription start site by examining FANTOM CAGE-seq data in the UCSC genome browser (assembly GRCh38/hg38).

We extracted aggregate data on ribosome-protected fragments for the same region of the spliced C9ORF72 Variant 2 mRNA from Ribocrypt (https://ribocrypt.org/). The database includes data from 393 experiments, with >1,000 samples.

Translation initiation probabilities at the main translon, the upstream A74 and A88 AUGs, and the near-cognate start codons were estimated with TISpredictor (https://www.tispredictor.com/kss). All predicted TISpredictor values are provided in **Supplementary Data S1**.

We predicted RNA secondary structures for the wildtype and for each mutant 5′ UTR with both the RNAfold Web Server (http://rna.tbi.univie.ac.at/) and RNAstructure server (https://rna.urmc.rochester.edu/RNAstructure.html). For the modeling, the stem of each stem-loop was operationally defined as the set of nucleotides with > 0.99 base-pairing probability under the RNAstructure prediction of the wild-type sequence. The predicted free energy of SL1, SL2 and SL3 was obtained from the RNAfold prediction; log base 2 transformed per-element ΔΔG values relative to wild-type were used as element-wise scores in the modeling.

We obtained the evolutionary conservation scores across 100 vertebrates from the UCSC Genome Browser (PhyloP 100-way; Phylip100).

RNA modifications were predicted using (http://www.rnanut.net/rnam5cfinder/, and internal ribosome entry sites were screened with IRESpy (https://irespy.shinyapps.io/IRESpy/). Neither provided positive results.

### SHAPE chemical probing of the V2 5′ UTR

Templates for *in vitro* transcription were generated by PCR from a psiCHECK-2 plasmid carrying the sequence of interest, using a forward primer that introduced the T7 promoter as a 5′ overhang together with the SHAPE-cassette sequences flanking the insert (5′ cassette: 5′-GGCCTTCGGGCCAA-3′; 3′ cassette: 5′-GAACCGGACCGAAGCCCGATTTGGATCCGGCGAACCGGATCGA-3′), as previously described ^20^. PCR was performed with Phusion polymerase (New England Biolabs) according to the manufacturer’s instructions. For each construct, four parallel 50 µL transcription reactions were prepared using in-house T7 RNA polymerase at MgCl₂ concentrations of 20, 40, 60 and 80 mM (a standard MgCl₂ screen), and the optimal MgCl₂ concentration (40 mM) was used to produce RNA for chemical probing experiments. Each 50 µL reaction contained 8 µL of PCR-amplified template, the indicated MgCl₂, 8 mM rNTPs, 5 % (w/v) PEG 8000, 1× transcription buffer (5 mM Tris-HCl pH 8.0, 5 mM spermidine, 10 mM DTT) and 6 µg T7 polymerase, incubated at 37 °C for 3 h. Transcribed RNAs were purified by ethanol precipitation and resuspended in nuclease-free water.

We performed Selective 2′-hydroxyl acylation analysed by primer extension (SHAPE) probing as described, screening multiple concentrations of probe to confirm single-hit kinetics ^49^. Cassette-extended RNAs containing the universal primer-binding site were diluted to 5 pmol, snap-cooled (95 °C, 3 min; ice, 10 min) and refolded at 37 °C for 20 min in folding buffer (100 mM NaCl, 100 mM HEPES pH 8.0, 10 mM MgCl₂). Folded RNAs were treated with either 1 mM, 6 mM or 10 mM 1-methyl-7-nitroisatoic anhydride (1M7) for 15 min at 25 °C; an untreated control was processed in parallel with DMSO in place of 1M7. Modified and untreated RNAs were purified using RNAClean XP beads (Beckman Coulter), reverse-transcribed with SuperScript IV (Invitrogen), and the cDNAs were amplified by PCR with PrimeStar GXL polymerase (Takara Bio) supplemented with 2 % DMSO. Double-stranded products were purified with AMPure XP beads (Beckman Coulter) and stored at −20 °C.

PCR products were end-repaired using the NEBNext Ultra II End Repair Module (New England Biolabs), barcoded with the Native Barcoding Kit 24 V14 (Oxford Nanopore Technologies) and ligated with motor protein according to the manufacturer’s instructions. Libraries were sequenced on a Flongle flow cell (FLO-FLG114, Oxford Nanopore Technologies, Oxford, UK). Reads were processed with ShapeMapper2 ^50^, and downstream analysis, including reactivity normalisation and SHAPE-constrained secondary-structure modeling was carried out with RNAvigate ^51^ using default parameters. Normalized SHAPE reactivities were used as constraints for RNA-structure modeling; nucleotides flagged with NaN reactivity (insufficient read depth) were treated as unconstrained. We conducted three independent replicates.

For visualisation of the wild-type V2 5′ UTR in **Figure 1d**, the SHAPE-derived secondary-structure model is the lowest free energy structural representation. For the mutational screen, where SHAPE was not feasible for every construct, the impact of each substitution on stem-loop stability was scored computationally as described above. SHAPE sequencing data is available upon request.

### Construction of reporter plasmids

We designed recombinant sequences to target sequence elements and structures as described in the text and in **Supplementary Data S1.** All dual-luciferase reporters were built on the psiCHECK-2 backbone (Promega), which encodes Renilla and Firefly luciferases and carries ampicillin resistance. DNA fragments were synthesized as gBlocks or Ultramer oligonucleotides (Integrated DNA Technologies) and inserted immediately upstream of the Renilla luciferase coding sequence at the NheI site by Gibson Assembly (NEBuilder HiFi DNA Assembly Cloning Kit, New England Biolabs) or by restriction-ligation (New England Biolabs). C9ORF72 5’UTR reporters comprised the full-length wild-type 5’UTR and recombinant variants, including the editing reporter - a C9ORF72 V2 5’ UTR containing the target adenosines A74 and A88. Larger replacements carried repeats of an unstructured 14-nt sequence lacking predicted start codons or basepairing (M14: AACAGAACAGACAG)(**Supplementary Figure S3**).

Additional 5’UTR variants were generated by site-directed mutagenesis (Q5 Site-Directed Mutagenesis Kit, New England Biolabs; or QuikChange Multi Site-Directed Mutagenesis Kit, Agilent Technologies), with primers designed using NEBaseChanger (https://nebasechanger.neb.com/) or the QuikChange Primer Design tool (https://www.agilent.com/store/primerDesignProgram.jsp) and synthesized by Integrated DNA Technologies.

We generated six chimeric reporters by fusing C9ORF72 V2 5′ UTR nucleotides 1-51 to the last 44 nucleotides of ACTB 5′ UTR (**Supplementary Figure S8**). All six constructs carried six substitutions in the C9ORF72 part (at position 4, 14, 32, 36, 48, 49) that disabled the frame-0 uTranslon, removing initiation at F0. Three constructs retained the wild-type nucleotides at positions 8, and 41, restoring the F2 initiation module. Where indicated, a downstream cassette replaced ACTB nucleotides 23 - 28 with either a start-stop element identical in position and composition to that in wild-type C9ORF72 V2 or a u9 uTranslon. All sequences are provided in **Supplementary Data S1**. We synthesised inserts as gBlocks (Integrated DNA Technologies) with 24-bp and 22-bp homology arms to the NheI-linearised backbone and joined them by Gibson Assembly (NEBuilder HiFi DNA Assembly Kit, NEB).

Assembled plasmids were transformed into *E. coli* NEB Stable (C2987H, New England Biolabs), selected on LB-agar with 100 µg ml⁻¹ ampicillin, and cultured in LB-ampicillin. Plasmid DNA was purified (QIAprep Spin Miniprep Kit, QIAGEN) and inserts were verified by Sanger sequencing across the NheI junction in both directions (Azenta Life Sciences). Full construct and primer sequences are provided in **Supplementary Data S1**.

### In cellulo evaluation

We procured HEK293T cells from a commercial source (ATCC). These cells were maintained in a humidified incubator at 37 °C with 5% CO₂, cultured in Dulbecco’s Modified Eagle Medium (DMEM, Gibco) supplemented with 10% fetal bovine serum (FBS, Gibco, Corning). Cell passaging or transfection was performed when cultures reached 70-80% confluency to ensure optimal growth conditions. For each biological replicate, 1 ug of reporter plasmid was used for transfection. Prior to transfection, the plasmid was mixed with transfection reagents according to the manufacturer’s protocol, using P3000 enhancer (Invitrogen), Lipofectamine 3000 reagent (Invitrogen), and Opti-MEM reduced serum medium (Gibco) for 20 minutes at room temperature to facilitate efficient liposome formation. Following this incubation, the transfection mixture was added to the cells, which were then incubated overnight at 37°C. Cells were tested regularly for mycoplasma contamination and were consistently negative.

Stress was induced by treating the transfected HEK293T cells with 1 μM thapsigargin (Sigma-Aldrich) for 4 hours at 37°C in a humidified incubator with 5% CO₂. Following stress induction, cells were harvested for detection of stress markers, including ATF4 expression and eIF2α phosphorylation using Western Blot. Cell lysates were prepared in RIPA buffer supplemented with protease and phosphatase inhibitors (Thermo Fisher Scientific), separated by SDS-PAGE, transferred to PVDF membranes, and probed with primary antibodies against ATF4 (Invitrogen), phospho-eIF2α (Ser51) (Cell Signaling Technology), total eIF2α (Cell Signaling Technology) and α-tubulin (Cell Signaling Technology), followed by incubation with spectrally distinct near-infrared secondary antibodies: Goat Anti-rabbit 680RD (Li-Cor), Goat Anti-Mouse 800CW (Li-Cor) according to the manufacturer’s protocol.. We imaged the membranes on a LI-COR Odyssey imaging system and band intensities were quantified using ImageJ software. All experiments were performed with at least three biological replicates.

### Protein quantification using dual-luciferase assay and mutant screen

We systematically screened 76 different C9ORF72 V2 constructs for changes in reporter protein expression relative to wild-type. Mutants targeted all predicted sequence and structure elements and many of their combinations. All experiments were conducted in at least triplicate. RNA was quantified for select constructs. All data are reported in **Supplementary Data S1.**

At 24-36 hours post-transfection, we rinsed HEK293T cells with Dulbecco’s Phosphate-Buffered Saline (DPBS, Gibco) to remove residual media, lysed them in 1X Passive Lysis Buffer (Promega Corporation) for 20-30 minutes at room temperature. For luminescence measurement, cell lysate was combined with LAR II reagent (Promega Corporation) to detect Firefly luciferase activity, followed by the addition of Stop & Glo reagent (Promega Corporation) to assess Renilla luciferase levels, all according to the manufacturer’s protocol. We quantified luminescence on a Tecan Spark Microplate Reader, and expressed reporter activity as the Renilla/Firefly ratio, normalized to the wild-type construct within each biological replicate.

Editing-dependent reporter activity was measured at 48 hours after ASO transfection and was expressed as the Renilla/Firefly ratio, normalized to the matched no-guide control within each biological replicate.

### RNA extraction, RT-qPCR and reverse transcription

We harvested transfected HEK293T cells by resuspension in DPBS (Gibco) and centrifugation, then extracted total RNA with the RNeasy Mini Kit (QIAGEN). For the mutant-screen RT-qPCR we further purified the RNA with the RNA Clean and Concentrator-5 kit (Zymo Research). Purified RNA was treated with DNase I to remove residual reporter plasmid, using paired +RT and −RT controls.

We quantified reporter-transcript levels by one-step RT-qPCR (Luna Universal One-Step RT-qPCR Kit, New England Biolabs) with Firefly- or Renilla-specific primers in 10 µL reactions on a QuantStudio 6 Flex Real-Time PCR System (Applied Biosystems, 384-well format). We normalized Renilla Ct to Firefly Ct (ΔCt) and calculated fold change relative to the wild-type construct (mutant screen) or the matched no-guide control (RNA editing) within each biological replicate.

For amplicon sequencing, first-strand cDNA was primed gene-specifically with the Renilla-RT primer (**Supplementary Data S1**), which anneals within the Renilla coding sequence downstream of the 5′UTR, selecting reporter transcripts spanning the 5′UTR-Renilla junction and excluding endogenous C9ORF72 mRNA.

### Antisense-oligonucleotide guided RNA editing

We used antisense oligonucleotides (ASOs) of 30-36 nucleotides length and antisense to the target region in the C9ORF72 5′UTR, with alternating 2′-O-methyl/2′-fluoro wings, phosphorothioate termini and a phosphodiester core, and a central 2′-deoxy triplet (dA–dN3U–dT) placing a 2’-deoxyisouridine mismatch opposite the target adenosine (guide chemistry adapted from ^52,53^). Single-site ASOs (30-mer) targeted A74 or A88 individually; a 36-mer targeted both simultaneously. For controls, we included a chemistry-matched scrambled ASO (non-targeting control) and an ACTB-targeting ASO (UAG→UGG; ADAR-activity positive control). Sequences and per-position chemistry are provided in **Supplementary Data S1**.

For editing experiments, ADAR1 p150 was supplied in trans from pmGFP-ADAR1-p150 (Addgene #117927). The psiCHECK-2 reporter containing the target 5’ UTR and ADAR1 p150 plasmids (1.25 µg total) were transfected with Lipofectamine 3000 in Opti-MEM; medium was refreshed ∼16 h later. Cells were re-seeded into 96-well plates (2 × 10⁴ cells per well, antibiotic-free medium) and ASOs were transfected with Lipofectamine RNAiMAX (0.2 µL per well). Targeted ASOs were transfected at final concentrations of 0, 0.3 or 1 µM; scramble controls at 1 µM. We conducted four independent replicates.

We generated amplicons spanning the two editing sites in C9ORF72 Variant 2 5’UTR, A74 and A88, together with ACTB positive control sites and submitted them to Azenta Amplicon EZ for library preparation and sequencing on an Illumina Miseq platform in 2 x 250 nts paired-end. We designed the primers and prepared the input according to Amplicon EZ protocol.

Azenta’s Genotyper processed the demultiplexed reads by aligning each read to the amplicon reference and reported per-position base counts for A, C, G and T, together with the total read depth for each sample (**Supplementary Figure S10**).

We calculated editing efficiency at each target adenosine from these counts A74 and A88 as G/(A+G) × 100 (inosine read as guanosine) and subtracted the matched control (no-guide or scramble) at each position for background correction. A position-wise scan quantified bystander editing.

All data for the RNA editing analysis are deposited in the NCBI Sequence Read Archive under BioProject accession PRJNA1514796. All analysis scripts are available at https://github.com/solomonhaizel/c9orf72-5utr-editing.

### Quantitative modeling

For the wild-type and each C9ORF72 mutant sequence, we assembled information on: the average log base 2 fold change in luciferase signal as measured in the mutant screen (mutant vs. wild-type, denoted Log2T), log base 2 fold change of the Kozak score as predicted by TISpredictor (see above), presence/absence of stop codons, log base 2 fold change of the free energy for SL1, SL2, and SL3. For the loss of a stop codon, we assumed ‘dummy’ values of the stop codon present (0.8) and absent (0.2). As TISpredictor does not score codon identity (e.g. canonical vs. near-cognate start codons), we weighted all scores with respect to start codon type. The scaling factor (weight) was 1 for AUG, 0.90 for CUG, GUG, UUG, and 0.60 for all other near-cognate start codons. Absent TISpredictor values were replaced by 0.2 and scaled as well. TISpredictor identified 16 near-cognate start codons in the wild-type and 6 new near-cognate start codons across the mutant sequences. All data are provided in **Supplementary Data S2**.

To reduce the number of variables, we merged the log base 2 TISpredictor fold-changes for all start codons in the same frame and upstream of the same in-frame stop codon into one frame (F0, F1, F2). We kept the start-stop (A74) and uTranslon (A88) separate. We then removed the data for the remaining downstream start codons and the main translon’s start codon due to their low information content (U83, G93, A96). We then removed the entries for the frame 2 stop codon (S77) as its presence/absence had little information content and was largely inconsequential for F2. This removal did not affect the way the start-stop was encoded. We then examined all main variables for cross-correlation to remove those with Pearson’s |R| > 0.80. There were none. However, we dropped SL3 from the data matrix as its information was entirely captured by the results from the start-stop and uTranslon mutants. Dropping SL3 was a practical choice - it reduces the number of variables while still retaining the same information. We also tested other ways of merging variables, e.g. merging A88 with F1, but found no modeling improvement (*not shown*).

The final data matrix contained the information for eight individual variables (main effects) for 76 independent observations (constructs): F0, F1, F2, SL1, SL2, S72 (stop codon in frame 0), A74 (start-stop), and A88 (uTranslon). **Supplementary Data S1** contains the data. Python scripts were developed with the support of generative AI tools.

We modeled Log2T using a hierarchical linear modeling framework that combined ordinary least squares (OLS) of the main effects with Elastic Net modeling of selected pairwise interactions on the residuals (after modeling the mains). After some initial optimization, we fixed the Elastic Net parameters at α = 0.3 and l1_ratio = 0.05. We used scikit-learn ElasticNet with max_iter = 100000, tol = 1 × 10⁻6, and cyclic coordinate selection. The intercept was not penalized.

For every training split, we standardized (normalized) each main predictor using the training-set mean and standard deviation and constructed all pairwise products of the standardized predictors. We then standardized interaction terms (Z-score normalization) using parameters estimated from the same training data. We excluded predictors or interactions with zero variance within a split and did not use held-out observations to estimate scaling parameters. This procedure applied to all cross-validation folds and bootstrap resamples.

We first screened interactions for the stability of their contribution to the model fit using leave-one-out cross-validation (LOOCV). Within each training fold, we fitted an OLS model to the standardized main effects (H1) and calculated its residuals. For each interaction, we defined the interaction stability score (ISS) as the absolute Pearson correlation between the standardized interaction and the H1 residuals. We then fitted an Elastic Net model (H2) to all interactions to predict these residuals and recorded whether each interaction had a nonzero coefficient and, when nonzero, its sign. Across LOOCV folds, we recorded the mean and standard deviation of the ISS, nonzero frequency, and sign stability.

We defined a global stable interaction pool by retaining interactions with nonzero frequency ≥0.60 and sign stability ≥0.80 and ranked retained terms by decreasing mean ISS. We removed interactions with an absolute Pearson correlation >0.80 with a main predictor or with a higher-ranked retained interaction. We then evaluated models containing the top K retained interactions, including K = 0, by LOOCV with fixed Elastic Net parameters listed above. For each K, predictions combined the H1 main-effect prediction with the H2 residual prediction; when K = 0, predictions used H1 alone. We selected the smallest K whose pooled out-of-fold R² was within 0.01 of the best observed value and designated the resulting interaction set as I_fixed (**Supplementary Data S2**).

We evaluated I_fixed by outer LOOCV. In each outer fold, we fitted H1 and H2 on the training observations and predicted the held-out observation. We calculated pooled out-of-fold R² as

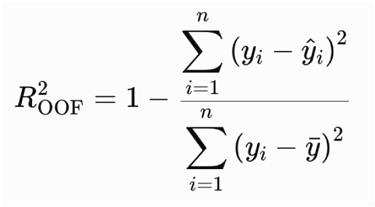

where y_bar is the mean of observed y (Log2T) across all n samples. We used one held-out prediction per observation rather than averaging fold-specific R² values.

Prior to final pruning, refinements, and model interpretation, we fitted the I_fixed selected interactions and mains with a direct Elastic Net model containing all main effects and I_fixed, evaluated using pooled LOOCV R², and repeated the 100-replicate conditional bootstrap for the same predictor set. We recalculated scaling within each bootstrap sample and recorded standardized coefficients with respect to their mean, standard deviations, and 2.5th and 97.5th percentiles.

Finally, we manually pruned the model by testing if removal of interactions with very low absolute coefficients (<0.05) improved LOOCV R². In addition, we calculated the cross-correlation matrix for all main variables and all interactions and selected for each interaction the set of interactions with Pearson’s |R| > 0.80 (**Supplementary Data S2**). We manually screened these interactions for better interpretability than the originally selected interaction and substituted with the new interaction if it resulted in the same LOOCV R². We then optimized α and l1_ratio and ran the final Elastic Net model with all mains and five interactions (two of which substituted) with α = 0.01 and l1_ratio = 0.01.

**Supplementary Data S2** contains output from the major modeling steps and the final predictions. Code development was supported by AI tools. All code and corresponding pseudocode is available at https://github.com/cvogelnyu/C9ORF72-translation-model.

### Split plot analysis

For the split plot analyses, we used the assembled data and defined a main and modifier variable from the interaction. We then split the data into two equal parts depending on high and low values for the modifier (n=38 each). For each part separately, we regressed the main variable with the measured log base 2 translation change and noted the slope and R^2^ of the regression (**Supplementary Data S2**). Python code to perform the split plot analysis is provided at https://github.com/cvogelnyu/C9ORF72-translation-model.

### Other bioinformatics and statistics methods

To re-analyze published transcriptomics data, we downloaded cerebellar RNA-seq from (GEO (GSE67196) from controls (n = 8), and C9-ALS patients (n = 8) and aligned the data to GRCh38 with STAR v2.7.11b (GENCODE v45, --sjdbOverhang 100). We obtained gene-level counts with --quantMode GeneCounts (reverse-stranded libraries). Total C9ORF72 (ENSG00000147894, all variants) counts were normalized using DESeq2 median-of-ratios size factors and compared by two-sided Mann-Whitney U test. Statistical tests were performed in SciPy 1.15.3 (Python 3.10.12); figures were generated in R with ggplot2. All scripts are provided at https://github.com/solomonhaizel/c9orf72-cerebellum-reanalysis.

## Supplementary Files

### Supplementary Information

This file contains Supplementary Figures/Tables.

### Supplementary Data File S1

This file contains additional experimental information.

### Supplementary Data File S2

This file contains additional information for the computational analysis.

## Acknowledgements

The authors acknowledge funding by the National Institutes of Health (R35GM127089 to C.V.; T32AG052909 to S.H.) and the Zegar Family Foundation for their generous support. A.N.J. acknowledges funding of NIH 1R35GM160278 (MIRA) and NSF grant 224667. L.S.V. was supported by the Czech Science Foundation (grant 26-21035S), and ERDF and MEYS (CZ.02.01.01/00/22_008/0004575 RNA for therapy).

## Author contributions

S.H. and C.V. conceived the project, conceptualized the experiments, analyzed the data, and wrote the manuscript. S.H. lead all experimental efforts. A.V.C. performed RNA SHAPE probing and analysis, and edited the manuscript. A.N.K, L,L and L.M cloned, purified, and performed experiments and edited the manuscript. R.A, S.Kher, S.Khazhinsky, X.G, H.Z, C. C, A.W, J.J-B, and S.K. assisted with cloning the various constructs.

